# Formic acid modulates latency and accuracy of nestmate recognition in carpenter ants

**DOI:** 10.1101/2021.04.28.441775

**Authors:** David Baracchi, Martin Giurfa, Patrizia d’Ettorre

## Abstract

Decision-making processes face the dilemma of being accurate or faster, a phenomenon that has been described as speed-accuracy trade-off (SAT) in numerous studies on animal behaviour. In social insects, discriminating between colony members and aliens is subjected to this trade-off as rapid and accurate rejection of enemies is of primary importance for the maintenance and ecological success of insect societies. Recognition cues distinguishing aliens from nestmates are embedded in the cuticular hydrocarbon (CHC) layer and vary among colonies. In walking carpenter ants, exposure to formic acid (FA), an alarm pheromone, improves accuracy of nestmate recognition by decreasing both alien acceptance and nestmate rejection. Here we studied the effect of FA exposure on the spontaneous aggressive mandible opening response of harnessed *Camponotus aethiops* ants presented with either nestmate or alien CHCs. FA modulated both MOR accuracy and the latency to respond to odours of conspecifics. In particular, FA decreased MOR towards nestmates but increased it towards aliens. Furthermore, FA decreased MOR latency towards aliens but not towards nestmates. As response latency can be used as a proxy of response speed, we conclude that contrary to the prediction of the SAT theory, ants did not trade off speed against accuracy in the process of nestmate recognition.

**Summary statement:** Exposure to an alarm pheromone increases both latency and accuracy of the response to recognition cues in ants

## Introduction

The recognition of group members is important for the evolution of cooperation and the maintenance of social life (Hamilton 1987). In social insects, discriminating colony members from aliens allows to direct appropriately altruistic behaviours without incurring in the cost of cooperating with intruders. Moreover, recognizing and reacting promptly to social parasites, robbers or predators is vital for colony success. As a result, insects living in social groups typically excel in discriminating friends and foes (d’Ettorre and Lenoir 2010). Social recognition systems are based on a multitude of cues from different sensory modalities among which vision and olfaction play a significant role (Tibbetts 2002; van Zweden and d’Ettorre 2010; Baracchi et al. 2016). In ants, recognition systems are predominantly based on the layer of hydrocarbons coating the cuticle of individuals, which defines the chemical signature of colonies (d’Ettorre and Lenoir 2010; Bos and d’Ettorre 2012). Cuticular hydrocarbons (CHCs) constitute a blend of many chemical compounds, mainly linear alkanes, alkenes and methyl-branched alkanes (van Zweden and d’Ettorre 2010), which vary qualitatively among different species, and quantitatively among colonies of the same species, or even among individuals belonging to different morphological or physiological castes (Vander Meer and Morel 1998; Monnin 2006). The sophisticated olfactory system of ants detects CHCs at very short distance (Brandstaetter et al. 2008) and resolves up to the individual level the identity of opponent ants (D’Ettorre and Heinze 2005), securing the nest from exploiters.

CHCs are not the only chemical cues that mediate social interactions in ants and other social insects. Volatile pheromones are also used to alert colony members to coordinate their defence against exploiters (Blum 1969; Nouvian et al. 2016). Pheromones are intraspecific chemical messengers that trigger context and signal-specific, adaptive responses (Karlson and Lüscher 1959; Wyatt 2014). Their primary function is to convey a message to one or more receivers, eliciting thereby a fast, highly predictable, and adaptive response. Yet, pheromones are not just chemical messengers. Recent work has uncovered a novel function for these substances, namely the modulation of the subjective evaluation of reinforcing stimuli (e.g. reward or punishment). Pheromones can thus modify the responsiveness to aversive or appetitive stimuli (Urlacher et al. 2010; Baracchi et al. 2017; Rossi et al. 2018; Baracchi et al. 2020). Such a modulatory effect was also detected when *Camponotus* ants were pre-exposed to the alarm pheromone formic acid (FA) in the context of social interactions (Rossi et al. 2019). In this case, pheromone pre-exposure improved nestmate discrimination accuracy by increasing aggressive behaviours towards aliens, while decreasing simultaneously aggression erroneously directed towards nestmates (Rossi et al. 2019). Although the exact neural mechanisms underlying this modulatory action of FA remain to be elucidated, it was suggested that this pheromone modulates attentional processes and thus the sensitivity to recognition cues (Rossi et al. 2019).

Although accuracy is certainly a crucial aspect of any recognition process, the speed of recognition is equally important (Heitz 2014). Both variables are intimately connected in many decision-making processes (Wickelgren 1977; Heitz 2014) and may operate in orthogonal ways as decision accuracy depends on being well informed, which requires time. On the contrary, being faster in a decision process could occur at the expense of being accurate. The relationship between speed and accuracy in decision making has been refereed as the speed–accuracy trade-off (SAT) (Busemeyer and Townsend 1993). Examples of SAT have been described for several social insect species, in many different ecologically relevant tasks, including foraging, predator detection, prey choice and communication (Wickelgren 1977; Franks et al. 2003; Ings and Chittka 2008; Trimmer et al. 2008; Chittka et al. 2009). For instance, when foraging bumblebees were tested in a colour discrimination task, some bees made rapid choices but with low precision, while other bees were slower but highly accurate (Chittka et al. 2003).

Here we focused on nestmate recognition in carpenter ants and studied whether the alarm pheromone FA affects not only the accuracy (Rossi et al. 2019) but also the speed of this process in an attempt to determine to what extent pheromones act on SAT processes. To this end, we pre-exposed individually harnessed ants to FA and quantified afterwards their mandible opening response (MOR) to a glass rod coated with alien or nestmate CHCs (Guerrieri and d’Ettorre 2008). This stereotyped defensive response has been already used to study both within- and between-species aggression in various ant species and aversive associative learning of carpenter ants (Desmedt et al. 2017). We thus determined if the speed and the accuracy of the recognition process are traded off and affected by pheromone pre-exposure.

## Material and Methods

### Study Species and Housing

We used four queen-right colonies of *Camponotus aethiops* (Latreille 1798) collected in 2016 at Pompertuzat (Midi-Pyrénées, France). Colonies were kept under controlled laboratory conditions (25°C, light-dark cycle = 12:12, ∼ 50% relative humidity). Each colony was housed in a plastic box (26 × 19 × 10 cm) with plaster floor connected by a tube to another box conceived as a foraging arena (26 × 19 × 10 cm) containing sand on the floor. The nest box was covered by cardboard in order to make it dark while the foraging arena was exposed to light. The inner faces of the two boxes were coated with Fluon® (AGC Chemicals Europe, Thornton Cleveleys, Lancashire, UK) to prevent ants from escaping. Ants were fed twice a week with pieces of crickets and flour worms for proteins and honey/apple mix for carbohydrates and vitamins. Water was provided *ad libitum*.

### Nestmate recognition assay

We designed an experiment to determine whether the alarm pheromone formic acid (FA, analytical grade, Sigma-Aldrich) modulates the latency and the accuracy of the responsiveness to nestmate and non-nestmate odours. On each experimental day, medium size forager ants were gently collected with tiny forceps from the foraging arena of one of the four colonies. Ants were immediately cold anesthetized on crushed ice for a few minutes and individually harnessed in small plastic holders. A small strip of adhesive tape between the head and the thorax was used to immobilize the ants, so that they could only freely move their antennae and mouthparts (Guerrieri and d’Ettorre 2008). Once harnessed, ants were kept resting for three hours in a dark and humid place at room temperature (about 60-70% relative humidity, 24 ± 2°C) to let them acclimatize to the new restraining situation. After resting, ants were randomly allocated to either the control group and exposed to pure water (solvent) or to the experimental group and exposed to FA.

A first assay (test 1) was performed before exposure to quantify basal responsiveness to nestmate and non-nestmate CHCs. Ant responsiveness to these chemicals was quantified using the mandible opening response (MOR), (Guerrieri and d’Ettorre 2008). The test entailed eight presentation trials: four nestmate trials (A) and four non-nestmate trials (B) in a pseudorandom sequence, such as ABABBABA, so that the same stimulus (A or B) was never presented more than twice consecutively. A 12-minute inter-trial interval was used. During each trial, one ant at a time was placed under a stereomicroscope (Leica S8 APO, magnification 10 ×) in order to better visualize its MOR. Each trial lasted 25 s and consisted of 10 s of familiarization with the experimental context, 10s of stimulus (nestmate or non-nestmate CHCs) presentation and 5 s of post-stimulus resting in the setup. Each chemical stimulus was presented to the harnessed ant on a glass rod whose tip was previously coated with the CHC extract of either nestmate or non-nestmate ants (see below). The glass rod was carefully manoeuvred by means of a micromanipulator (WPI, M33) to avoid contaminations. Upon stimulation, the rod was placed always at the same distance (2 mm from the head) of the antennae. Each stimulus was preceded by the presentation of a clean rod (presented by hands) in order to familiarize the ants with the visual component of this stimulus.

Fifteen min after the end of this first assay (test 1), ants were exposed either to FA (experimental group) or to the solvent alone (pure water, control group) to determine if pheromonal exposure modified their MOR responses. To this end, harnessed ants were individually confined for 15 min in a 50 ml plastic bottle containing a filter paper (1 x 5 cm) soaked either with the pheromone or pure water (Rossi et al. 2019). The entire procedure was performed under a hood. FA was diluted to 12% (3 μl pheromone + 22 μl water, equivalent to one third of the content of one poison gland (Stumper 1952). Control ants were exposed to 25 µl of water. After exposure, ants were kept resting for additional 30 min (Rossi et al. 2019) and then tested again (test 2) for responsiveness to nestmate and non-nestmate odours using the same procedure as in test 1.

CHC extracts were obtained by washing pools of 5 nestmate or non-nestmate ants in 2.5 ml of solvent (pentane, HPLC grade, Sigma Aldrich) for 10 min (Rossi et al. 2019). The amount of nestmate and non-nestmate odour used in each presentation was equivalent to that of a single ant. The tips of the rods were coated by adding drops of the chemical extracts using a micropipette and the rods were let dry for 1 h before starting the experiment to ensure that the solvent (pentane) evaporated. To avoid real replicates during the eight presentation trials within each assay (test 1 and test 2), alien and nestmate extracts were obtained from 4 pools of alien and nestmate ants, respectively. In the case of non-nestmates, each pool belonged to a different colony. Each presentation was video recorded from above with an integrated microscope camera. The latency to display the MOR from the moment in which the rod was positioned at 2 mm from the head and the occurrence of MOR (yes/no) to each stimulus presentation were quantified.

### Locomotor activity assay

In order to determine whether FA merely affected motor responses, thus influencing the observed MOR results, we designed a simple assay to monitor the locomotor activity of free walking ants, pre-exposed either to FA or to water, which is described in the Supplementary Materials. The results show that FA did neither impair nor modulate the locomotor activity of ants (Fig. S1).

### Data analysis and statistics

To study the effect of FA in terms of population response the proportion of reacting ants and the speed of their response to nestmate and non-nestmate stimulations, after and before exposure, were analysed using ANOVA designs. For testing accuracy, individual ant responses (MOR: 1 or 0) were examined using generalized linear mixed models (GLMMs) with a binomial error structure - logit-link function -, *glmer* function of R package *lme4* (Bates et al. 2014). The speed of the response was analysed using GLMMs fit with Poisson family distribution and identity link function. Q–Q plots and scatterplots of the residuals of the model were checked visually for normal distribution and homoscedasticity. In both cases, independent analyses were performed for ants exposed to water and ants exposed to FA. In all the models ‘*Ant response*” was entered as dependent variable, “*Treatment time*” (before/ after exposure to either water or FA) and “*Odour stimulus*” (nestmate/non-nestmate extract) as fixed factors, and “*Trial*” as covariate. Moreover, ‘*Individual identity*’ (*IDs*) was considered as a random factor to allow for repeated-measurement analysis. Colony of origin was also entered as random factor. When necessary, models where optimized with the iterative algorithms BOBYQA or Nelder-Mead. In each analysis, several models were run and compared to identify significant interactions between fixed factors and/or covariates and the significant model with the highest explanatory power (i.e. the lowest AIC value) was retained. Interactions, wherever significant, are reported in the text. Tukey’s post-hoc tests were used to detect differences between the different groups (*lsmeans* function from R package *lsmeans* (Lenth and Lenth 2018).

To study the effect of FA at the individual level, for each tethered ant we calculated a nestmate and a non-nestmate MOR Score (MS). The former was quantified as the sum of MORs to the four nestmate presentations while the latter was quantified as the sum of MORs to the four non-nestmate presentations. Thus, both MSs could range from 0 to 4. In the case of nestmates, higher MS values correspond to incorrect responses (i.e. aggressive display towards a nestmate). On the contrary, in the case of non-nestmates, higher MS values correspond to correct responses (i.e. aggressive display towards an alien). MSs were calculated for test 1 (before exposure) and for test 2 (after exposure) and compared by means of Wilcoxon signed-rank tests. We also calculated a Latency Score (LS) for each individual ant presented with nestmate and with non-nestmate CHCs. In the case of nestmates, the LS corresponded to the mean latency of aggressive responses upon the four nestmate odour presentations while in the case of non-nestmates, the LS corresponded to the mean latency of aggressive responses upon the four non-nestmate odour presentations. Both LSs were calculated for test 1 (before exposure) and for test 2 (after exposure) and compared by means of a Wilcoxon signed-rank test.

Finally, to test for the existence of a latency vs. accuracy trade-off in nestmate and non-nestmate recognition, Spearman rank tests were used to correlate MSs for nestmates and non-nestmates with nestmate and non-nestmate LSs respectively after and before FA/Water exposure. All statistical analyses were performed with R 4.0.3 (Team 2020).

## Results

In natural conditions, medium-size forager ants are typically aggressive towards alien ants and tolerant towards nestmates (Larsen et al. 2016). In the laboratory conditions in which the MOR bioassay was performed, harnessed ants reproduced this behaviour and displayed MOR to alien CHCs. In test 1 (before pheromone exposure), both FA and Water (control) groups, which were in principle identical at this point, reacted more aggressively towards the four non-nestmate presentations than to the four nestmate presentations (Fig. 1AB: within each ant category – “FA exposed” and “Water exposed” – compare the two proportions labelled as ‘Before’). This difference in the proportion of ants responding to either odour was significant (GLMM, Water group: *Odour stimulus*: χ^2^ = 5.63, df = 1, p = 0.018; FA group: *Odour stimulus*: χ^2^ = 5.93, df = 1, p = 0.015, Fig. 1AB and Fig. S2). Pheromone exposure induced a change in the proportion of ants responding with MOR to nestmate and non-nestmate odours (GLMM, *Odour stimulus * Treatment time*: χ^2^ = 36.35, df = 1, p < 0.0001, Fig. 1A and Fig. S2). In particular, FA exposure decreased erroneous MOR towards nestmates (GLMM, *Tukey post-hoc test*: Z = −6.05, p < 0.0001, Fig. 1A) while it increased correct MOR towards aliens, albeit in a non-significant way (GLMM, *Tukey post-hoc test*: Z = 2.32, p = 0.09, Fig. 1A). On the contrary, when ants were exposed to water, the proportion of individuals responding to nestmates and to aliens did not vary significantly (GLMM, *Odour stimulus * Treatment time*: χ^2^ = 1.21, df = 1, p = 0.27, Fig. 1B and Fig. S2).

**Figure 1:**
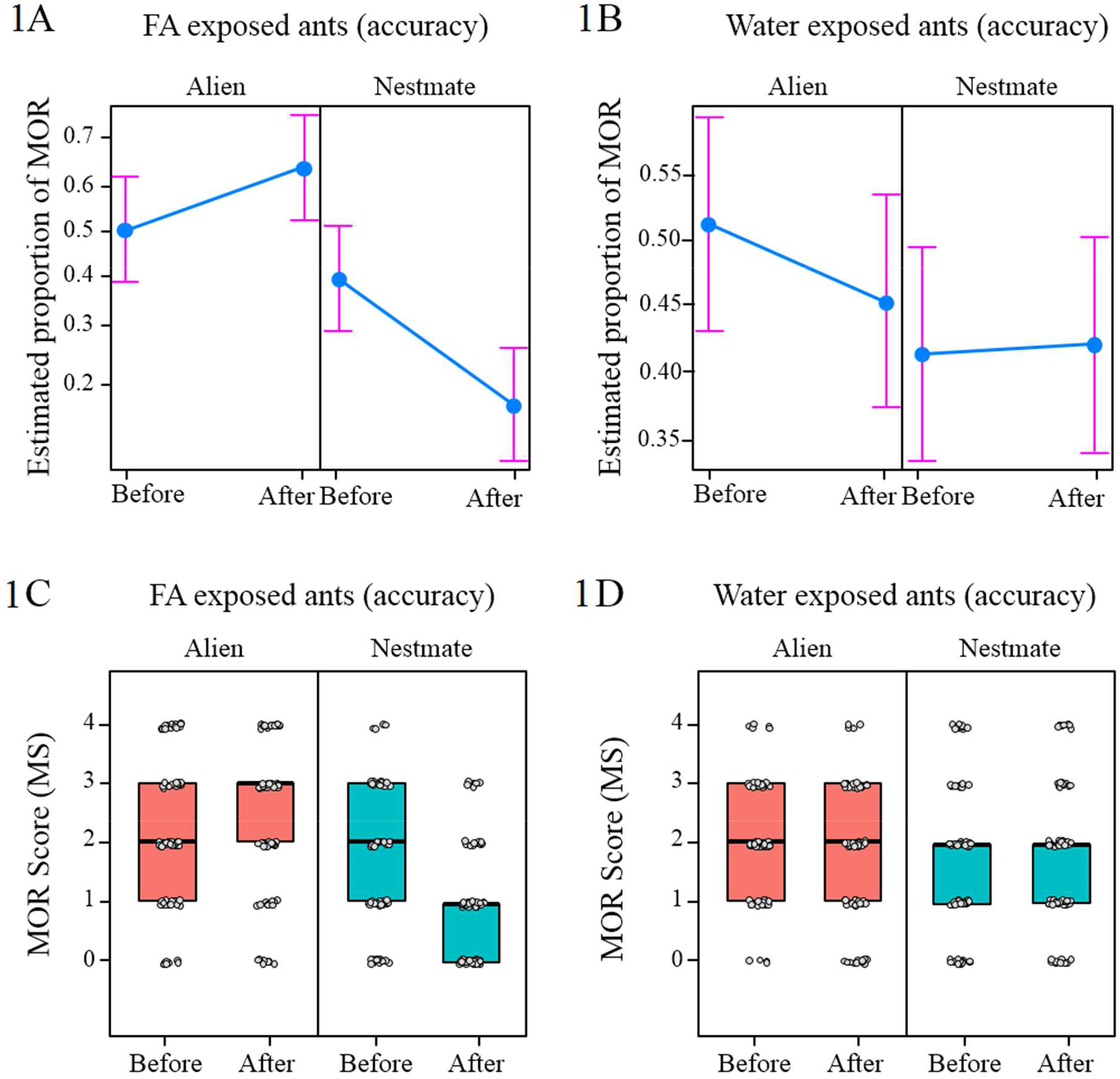
**(A-B)** Interaction plots of fitted means for the factors “*Treatment time*” (before/ after exposure to either FA or water) and “*Odour stimulus*” (nestmate/non-nestmate odours). **(A)** MOR was differently affected by FA exposure depending on the nature of the stimulus presented so that it increased towards non-nestmate odour (GLMM, *Tukey post-hoc test*: p = 0.09) and it decreased towards nestmate odour (GLMM, *Tukey post-hoc test*: p = 0.0001). **(B)** Ants exposed to water did not change their responsiveness neither towards nestmates nor to aliens (GLMM, *Odour stimulus * Treatment time*: p = 0.27. **(C-D)** Nestmate and non-nestmate MOR Score (MS) of individual tethered ants exposed either to FA or water. Boxes represent median, quartiles and max and min (upper and lower whiskers) MS values. Grey dots represent individual ants. **(C)** FA exposure tended to increase the MOR score (MS) to non-nestmates (p = 0.054) while it decreased it to nestmates (Wilcoxon test, p < 0.0001). **(D)** Water exposure did not affect the ants’ MS neither towards nestmates (p = 0.76) nor to aliens (p = 0.13).

In order to evaluate interindividual variability, we analysed responses in terms of individual MOR scores (MSs), which were computed both for responses to nestmate CHCs (i.e. the sum of responses to the four nestmate trials) and to non-nestmate CHCs (i.e. the sum of responses to the four non-nestmate trials). Fig. 1C shows that FA exposure significantly decreased responses to nestmates, (Wilcoxon test, n = 69, V = 108, p < 0.0001, Fig. 1C) and, to a lower extent, increased responses to non-nestmates, (Wilcoxon test: n = 69, V = 961, p = 0.054, Fig. 1C). On the contrary, exposure to water did not affect the individual MS, neither towards nestmates nor to aliens (Wilcoxon test, nestmates: n = 73, V = 805, p = 0.76; alien: n = 73, V = 572, p = 0.13, Fig. 1D).

At the population level, pheromone exposure induced a change in the mean latency of the MOR elicited by nestmate and non-nestmate odours (GLMM, *Odour stimulus * Treatment time*: χ^2^ = 555.4, df = 1, p < 0.0001, Fig. 2AB and Fig. S3). In particular, ants exposed to FA had a shorter MOR latencies towards alien CHCs (GLMM, *Tukey post-hoc* test: Z = −23.75, p < 0.0001, Fig. 2A) but did not change the MOR latency to nestmate CHCs (GLMM, *Tukey post-hoc test*: Z = −0.98, p = 0.76, Fig. 2A). Overall, MOR latency towards alien CHCs decreased over the presentations and tests following FA exposure (GLMM, *Trial*: χ^2^ = 146.3, df = 1, p < 0.0001). On the contrary, ants exposed to water did neither change the latency of MOR towards nestmate nor to alien CHCs (GLMM, *Treatment time*: χ^2^ = 0.17, df = 1, p = 0.68; *Odour stimulus* Treatment time*: χ^2^ = 1.01, df = 1, p = 0.31, Fig. 2B and Fig. S3). Overall, the MOR latency increased over the presentations (GLMM, *Trial*: χ^2^ = 86.1, df = 1, p < 0.0001).

**Figure 2:**
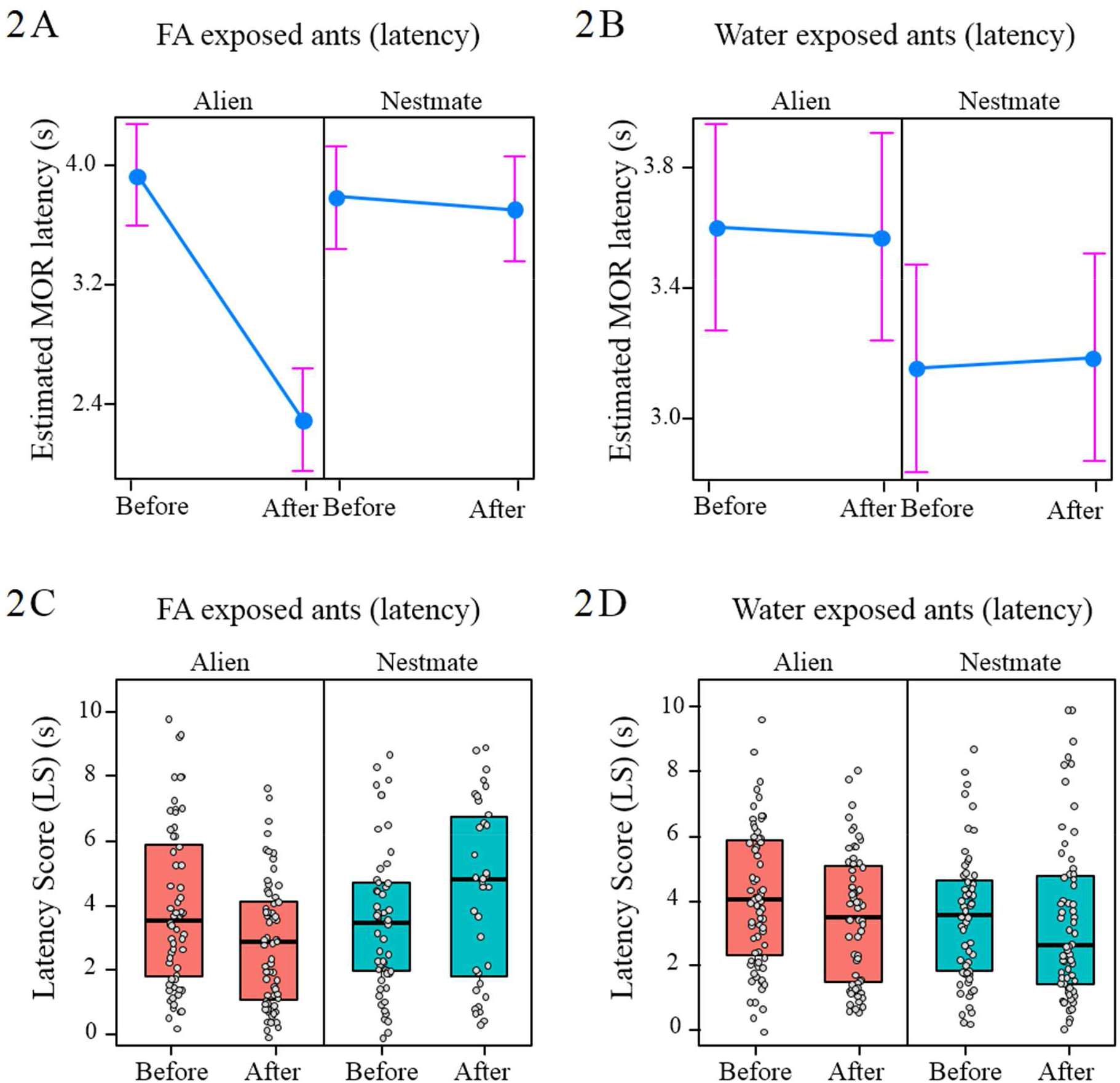
**(A-B)** Interaction plots of fitted means for the factors “*Treatment time*” (before/ after exposure to either FA or water) and “*Odour stimulus*” (nestmate/non-nestmate odours). **(A)** MOR latency was differently affected by FA exposure in a stimulus dependent manner so that it strongly decreased when ants were presented with non-nestmate odour (GLMM, *Tukey post-hoc test*: p < 0.0001) but it did not vary when the same ants were presented with nestmate odour (GLMM, *Tukey post-hoc test*: p = 0.76). **(B)** After water exposure ants did neither change the latency of MOR towards nestmates nor to aliens (GLMM, *Odour stimulus* Treatment time*: p = 0.31). **(C-D)** Nestmate and the non-nestmate latency Score (LS) of individual tethered ants exposed either to FA or water. Boxes represent median, quartiles and max and min (upper and lower whiskers) LS values. Grey dots represent individual ants. **(C)** FA exposure decreased the LS to non-nestmates (Wilcoxon test, p = 0.002) but not to nestmates (p = 0.12). **(D)** After water exposure ants slightly decreased LS towards aliens (p = 0.043, but not to nestmates (p = 0.37).

At the individual level, FA exposure significantly decreased the Latency Score (LS) to non-nestmates (Wilcoxon test: n = 56, V = 414, p = 0.002, Fig. 2C) but not to nestmates (Wilcoxon test: n = 31, V = 328, p = 0.12, Fig. 2C). After water exposure, ants slightly decreased their LS towards aliens (Wilcoxon test, n = 58, V = 593, p = 0.043, Fig. 2D), but not to nestmates (Wilcoxon test, n = 52, V = 566, p = 0.37, Fig. 2D).

We then combined our data about Latency Score (LS) and MOR Scores (MS) to analyse the existence of a speed versus accuracy trade-off. While the quantification of MS provides a measurement of response accuracy when the ants are confronted with alien or nestmate odours, the latency of their response informs about the potential speed of their response; typically shorter latencies are associated with faster responses and higher speed while longer latencies are associated with slower responses and slower speed.

We found that the response latency was not affected by the accuracy of both nestmate and non-nestmate odour recognition in ants before exposure. Precisely, individual nestmate MS did not correlate with nestmate LS (Spearman test, n = 113, ρ = 0.09, p = 0.35). Similarly, individual non-nestmate MS did not correlate with non-nestmate LS (Spearman test, n = 130, ρ = −0.17, p = 0.053). After exposure to FA or water, no significant correlation was found when we analysed the LSs and MSs of exposed ants for nestmates (FA: Spearman test, n = 35, ρ = −0.19, p = 0.28; Water: n = 63, ρ = −0.05, p = 0.68) and non-nestmates (FA: n = 64, ρ = −0.19, p = 0.14; Water: n = 62, ρ = 0.04, p = 0.77). Thus, ants did not trade-off these two aspects of the recognition process.

## Discussion

The ability to discriminate between nestmates and intruders allows colony cohesion and nest defence in social insects (Hamilton 1987). Accuracy is certainly a crucial aspect of the action component of the recognition process. Yet, the speed of the recognition is equally important (Heitz 2014). Therefore, while the existence of SAT necessarily imposes boundaries, the recognition process is expected to be as accurate and fast as possible (Wickelgren 1977; Heitz 2014). Over the course of evolution, the sensory systems of social insects have been strongly refined to achieve this goal (Stroeymeyt et al. 2010; van Zweden and d’Ettorre 2010; Ozaki and Hefetz 2014). Pheromones participate in this process as they have been naturally selected to facilitate communication and response coordination at the colony level (Blum 1969). Alarm pheromones, in particular, coordinate defensive responses of social groups and allow individuals to react promptly with stereotyped responses towards imminent dangers, such as the presence of enemies (Blum 1969; Nouvian et al. 2016).

In a previous study, we showed that FA, the alarm pheromone of several ant species, acts as cognitive modulator by enhancing nestmate discrimination in *Camponotus aethiops* ants, even when it is no longer present in the surroundings of the targeted ant (Rossi et al. 2019). In this case, FA exposure also increased aggressive behaviours of ants walking in an arena and confronted with aliens, while it decreased simultaneously erroneous aggression towards nestmates. Here we used a more controlled setup in which harnessed ants were exposed to CHCs of aliens or nestmates and confirmed our previous findings showing nestmate recognition improvement by FA in carpenter ants. In addition, we could evaluate the incidence of response latency in this process.

Our new results show that exposure to FA not only made ants more accurate in their aggressive responses but also modulated the latency of these responses. After FA exposure, those ants that still displayed erroneously MOR to nestmate odours, did it with the same latency. On the contrary, FA exposure reduced the latency of MOR towards non-nestmate odours. Thus, FA appears to act as a facilitator that speeds aggression towards the right targets. Most likely, these changes in response latency are relevant in natural scenarios, where faster attacks to non-nestmates would increase the probability of colony success.

The theory of SAT (Wickelgren 1977; Heitz 2014) predicts that correct decisions take longer while fast decisions are more error prone. Although we did not quantify response speed (i.e. the speed of a triggered MOR), we measured the latency of MOR, which can be used as a proxy of MOR speed (i.e. a shorter latency corresponds to a faster response consummation, while a longer latency corresponds to a slower response consummation). Our results show that carpenter ants did not trade off speed (latency) against accuracy, both before and after exposure to the alarm pheromone FA. The observed increased accuracy was not affected by the speed of the responses, as FA exposure enhanced both the accuracy and the latency of the responses. Although a trade-off between speed and accuracy has been described in various contexts involving decision making, and in different modalities (Chittka et al. 2009), there are cases in which a correlation between accuracy and sampling time has not been found both in insects (Ditzen et al. 2003) and in mammals facing olfactory-discrimination problems (Uchida and Mainen 2003). Notably, in a study on nestmate recognition by hover wasps, no SAT between speed and accuracy was found (Baracchi et al. 2015), suggesting that in this particular context SAT may be uncommon.

The increased accuracy in nestmate recognition induced by FA exposure may be explained by an enhanced sensitivity to CHCs (Rossi et al. 2019). It has been proposed that FA increases the amount of information (e.g., the number of detected CHCs) available to the ants, thus decreasing the perceived phenotypic overlap between nestmate and non-nestmate recognition cues (Rossi et al. 2019). Changes in recognition speed, which determined changes in response latency, cannot be explained by changes in motor abilities as the general locomotor activity of ants was unaffected by exposure to the pheromone (Supplementary Material). A possibility would be that FA affected attentional processes and enhanced motivation for the defensive task by acting on brain levels of neurotransmitters that have been associated with enhanced attention and aggressive responses. Attentional processes, similar to those described in vertebrates, have been characterized in insects both at the behavioral and neurobiological levels (Dyer and Chittka 2004; Giurfa 2004; Miller et al. 2011; van Swinderen 2011; Van Swinderen and Andretic 2011). In the fruit fly *Drosophila melanogaster*, visual attention for moving bars is mediated by a transient increase in a 20-30 Hz local field-potential recorded in a region of the brain called the medial protocerebrum (van Swinderen and Greenspan 2003). Current views relate dopamine levels in the insect brain with arousal levels (Van Swinderen and Andretic 2011). In consequence, attenuation of dopamine release in fly mutants attenuates the 20-30 Hz responsiveness to the visual object to be attended. On the contrary, pharmacological increase of dopamine rescues this responsiveness (Andretic et al. 2005). Thus, FA may upregulate dopamine levels in the brain, enhancing thereby attention in the context of nestmate discrimination. This hypothesis is sustained by findings on defensive responses in honey bees, which are triggered by the sting alarm pheromone component isoamyl acetate (IAA) (Nouvian et al. 2018). Exposure to IAA increases defensive responses and upregulates dopamine and serotonin levels in the bee brain. While serotonin has been directly related to aggressive responses in invertebrates (Kravitz 2000; Dierick and Greenspan 2007; Alekseyenko et al. 2010; Alekseyenko and Kravitz 2014; Alekseyenko et al. 2019), the dopamine component of the response may reflect the enhanced attention required to direct appropriately an attack that may have lethal consequences for the defender bee.

In conclusion, we found that FA improved nestmate recognition in *C. aethiops* by acting both on the accuracy (reducing erroneous responses) and on the latency of aggressive responses (reducing the latency of appropriate attacks). Our behavioural experiments do not allow identifying the mechanism of action of FA and neural analyses are necessary to determine if and how exposure to FA upregulates levels of biogenic amines that have been associated with aggressive responses and with attentional processes. Future research aimed at quantifying biogenic amine levels upon FA exposure and at specifically blocking/activating biogenic amines receptor might help to shed light on the underlying mechanisms of FA action. Our findings add to new perspectives developed recently positing that pheromone functions exceed the traditional framework of intraspecific communication for which they have been selected. Pheromones do more than conveying specific messages to members of the same species. In insects, for instance, they can modulate in the long term responsiveness to relevant stimuli (appetitive, aversive) in contexts that differ from the one for which the pheromone is used as a messenger (Baracchi et al. 2017; Rossi et al. 2018; Hostachy et al. 2019; Rossi et al. 2019; Baracchi et al. 2020; Murmu et al. 2020; Oberhauser et al. 2020; Rossi et al. 2020). Further studies are needed to clarify these novel functions of pheromones as neuromodulators and to understand their implications for the functioning of recognition systems in general.

## Author Contributions

DB and PdE conceived the study and designed the experiments. DB performed the experiments and carried out data analyses. DB, PdE and MG contributed to the writing of the manuscript.

## References

Alekseyenko OV, Chan YB, Okaty BW, Chang Y, Dymecki SM, Kravitz EA (2019) Serotonergic Modulation of Aggression in Drosophila Involves GABAergic and Cholinergic Opposing Pathways. Current Biology 29:2145-2156.e5

Alekseyenko OV, Kravitz EA (2014) Serotonin and the search for the anatomical substrate of aggression. Fly (Austin) 8:200–5

Alekseyenko OV, Lee C, Kravitz EA (2010) Targeted manipulation of serotonergic neurotransmission affects the escalation of aggression in adult male Drosophila melanogaster. PLoS One 5:e10806

Andretic R, van Swinderen B, Greenspan RJ (2005) Dopaminergic modulation of arousal in Drosophila. Current Biology 15:1165–1175

Baracchi D, Cabirol A, Devaud J-M, Haase A, d’Ettorre P, Giurfa M (2020) Pheromone components affect motivation and induce persistent modulation of associative learning and memory in honey bees. Communications biology 3:1–9

Baracchi D, Devaud J-M, d’Ettorre P, Giurfa M (2017) Pheromones modulate reward responsiveness and non-associative learning in honey bees. Scientific reports 7:1–9

Baracchi D, Petrocelli I, Chittka L, Ricciardi G, Turillazzi S (2015) Speed and accuracy in nest-mate recognition: a hover wasp prioritizes face recognition over colony odour cues to minimize intrusion by outsiders. Proceedings of the Royal Society B: Biological Sciences 282:20142750

Baracchi D, Turillazzi S, Chittka L (2016) Facial patterns in a tropical social wasp correlate with colony membership. The Science of Nature 103:80

Bates D, Mächler M, Bolker B, Walker S (2014) Fitting linear mixed-effects models using lme4. arXiv preprint arXiv:14065823

Blum MS (1969) Alarm pheromones. Annual Review of Entomology 14:57–80

Bos N, d’Ettorre P (2012) Recognition of social identity in ants. Frontiers in Psychology 3:83

Brandstaetter AS, Endler A, Kleineidam CJ (2008) Nestmate recognition in ants is possible without tactile interaction. Naturwissenschaften 95:601–608

Busemeyer JR, Townsend JT (1993) Decision field theory: a dynamic-cognitive approach to decision making in an uncertain environment. Psychological Review 100:432

Chittka L, Skorupski P, Raine NE (2009) Speed–accuracy tradeoffs in animal decision making. Trends in Ecology & Evolution 24:400–407

D’Ettorre P, Heinze J (2005) Individual recognition in ant queens. Current Biology 15:2170–2174

d’Ettorre P, Lenoir A (2010) Nestmate recognition. Ant ecology:194–209

Desmedt L, Baracchi D, Devaud J-M, Giurfa M, d’Ettorre P (2017) Aversive learning of odor–heat associations in ants. Journal of Experimental Biology 220:4661–4668

Dierick HA, Greenspan RJ (2007) Serotonin and neuropeptide F have opposite modulatory effects on fly aggression. Nature Genetics 39:678–82

Ditzen M, Evers J-F, Galizia CG (2003) Odor similarity does not influence the time needed for odor processing. Chemical Senses 28:781–789

Dyer AG, Chittka L (2004) Fine colour discrimination requires differential conditioning in bumblebees. Naturwissenschaften 91:224–227

Franks NR, Dornhaus A, Fitzsimmons JP, Stevens M (2003) Speed versus accuracy in collective decision making. Proceedings of the Royal Society of London Series B: Biological Sciences 270:2457–2463

Giurfa M (2004) Conditioning procedure and color discrimination in the honeybee Apis mellifera. Naturwissenschaften 91:228–231

Guerrieri FJ, d’Ettorre P (2008) The mandible opening response: quantifying aggression elicited by chemical cues in ants. Journal of Experimental Biology 211:1109–1113

Hamilton WD (1987) Discrimination nepotism: expectable, common, overlooked. Kin recognition in animals:417-437

Heitz RP (2014) The speed-accuracy tradeoff: history, physiology, methodology, and behavior. Frontiers in Neuroscience 8:150

Hostachy C, Couzi P, Portemer G, Hanafi-Portier M, Murmu M, Deisig N, Dacher M (2019) Exposure to conspecific and heterospecific sex-pheromones modulates gustatory habituation in the moth Agrotis ipsilon. Frontiers in Physiology 10

Ings TC, Chittka L (2008) Speed-accuracy tradeoffs and false alarms in bee responses to cryptic predators. Current Biology 18:1520–1524

Karlson P, Lüscher M (1959) ‘Pheromones’: a new term for a class of biologically active substances. Nature 183:55–56

Kravitz EA (2000) Serotonin and aggression: insights gained from a lobster model system and speculations on the role of amine neurons in a complex behavior. Journal of Comparative Physiology A 186:221–38

Larsen J, Nehring V, d’Ettorre P, Bos N (2016) Task specialization influences nestmate recognition ability in ants. Behavioral Ecology and Sociobiology 70:1433–1440

Lenth R, Lenth MR (2018) Package ‘lsmeans’. The American Statistician 34:216–221

Miller SM, Ngo TT, van Swinderen B (2011) Attentional switching in humans and flies: rivalry in large and miniature brains. Frontiers in Human Neuroscience 5:188

Monnin T (2006) Chemical recognition of reproductive status in social insects. In: Annales Zoologici Fennici. JSTOR, pp 515–530

Murmu MS, Hanoune J, Choi A, Bureau V, Renou M, Dacher M, Deisig N (2020) Modulatory effects of pheromone on olfactory learning and memory in moths. Journal of Insect Physiology 127:104159

Nouvian M, Mandal S, Jamme C, Claudianos C, d’Ettorre P, Reinhard J, Barron A, Giurfa M (2018) Cooperative defence operates by social modulation of biogenic amine levels in the honeybee brain. Proceedings of the Royal Society of London Series B: Biological Sciences 285:20172653.

Nouvian M, Reinhard J, Giurfa M (2016) The defensive response of the honeybee Apis mellifera. Journal of Experimental Biology 219:3505–3517

Oberhauser FB, Wendt S, Czaczkes TJ (2020) Trail pheromone does not modulate subjective reward evaluation in Lasius niger ants. Frontiers in Psychology 11:2515

Ozaki M, Hefetz A (2014) Neural mechanisms and information processing in recognition systems. Insects 5:722–741

Rossi N, Baracchi D, Giurfa M, d’Ettorre P (2019) Pheromone-Induced Accuracy of Nestmate Recognition in Carpenter Ants: Simultaneous Decrease in Type I and Type II Errors. The American Naturalist 193:267–278

Rossi N, d’Ettorre P, Giurfa M (2018) Pheromones modulate responsiveness to a noxious stimulus in honey bees. Journal of Experimental Biology 221

Rossi N, Pereyra M, Moauro MA, Giurfa M, d’Ettorre P, Josens R (2020) Trail pheromone modulates subjective reward evaluation in Argentine ants. Journal of Experimental Biology 223

Stroeymeyt N, Guerrieri FJ, van Zweden JS, d’Ettorre P (2010) Rapid decision-making with side-specific perceptual discrimination in ants. PLoS One 5

Stumper R (1952) Quantitative data on the secretion of formic acid by ants. Comptes rendus hebdomadaires des seances de l’Academie des sciences 234:149

Team R (2020) Core (2020). R: A language and environment for statistical computing R Foundation for Statistical Computing, Vienna, Austria URL https://www.R-project org

Tibbetts EA (2002) Visual signals of individual identity in the wasp Polistes fuscatus. Proceedings of the Royal Society of London Series B: Biological Sciences 269:1423–1428

Trimmer PC, Houston AI, Marshall JA, Bogacz R, Paul ES, Mendl MT, McNamara JM (2008) Mammalian choices: combining fast-but-inaccurate and slow-but-accurate decision-making systems. Proceedings of the Royal Society B: Biological Sciences 275:2353–2361

Uchida N, Mainen ZF (2003) Speed and accuracy of olfactory discrimination in the rat. Nature Neuroscience 6:1224–1229

Urlacher E, Francés B, Giurfa M, Devaud J-M (2010) An alarm pheromone modulates appetitive olfactory learning in the honeybee (Apis mellifera). Frontiers in Behavioral Neuroscience 4:157

van Swinderen B (2011) Attention in Drosophila. International Review of Neurobiology 99:51–85

van Swinderen B, Andretic R (2011) Dopamine in Drosophila: setting arousal thresholds in a miniature brain. Proceedings of the Royal Society B: Biological Sciences 278:906–913

van Swinderen B, Greenspan RJ (2003) Salience modulates 20-30 Hz brain activity in Drosophila. Nature Neuroscience 6:579–586

van Zweden JS, d’Ettorre P (2010) Nestmate recognition in social insects and the role of hydrocarbons. Insect hydrocarbons: biology, biochemistry and chemical ecology 11:222–243

Vander Meer RK, Morel L (1998) Nestmate recognition in ants. Pheromone communication in social insects 79

Wickelgren WA (1977) Speed-accuracy tradeoff and information processing dynamics. Acta Psychologica 41:67–85

Wyatt TD (2014) Pheromones and animal behavior: chemical signals and signatures. Cambridge University Press

